# Effect of Nrf2 Loss on Senescence and Cognition of Tau-Based P301S Mice

**DOI:** 10.1101/2022.10.31.514571

**Authors:** R. Riordan, W. Rong, Z. Yu, G. Ross, J. Valerio, J. Dimas-Muñoz, V. Heredia, K. Magnusson, V. Galvan, V.I. Perez

## Abstract

Cellular senescence may contribute to chronic inflammation involved in the progression of age-related diseases such as Alzheimer’s Disease (AD), and its removal prevents cognitive impairment in a model of tauopathy. Nrf2, the major transcription factor for damage response pathways and regulators of inflammation, declines with age. Our previous work showed that silencing Nrf2 gives rise to premature senescence in cells and mice. Others have shown that Nrf2 ablation can exacerbate cognitive phenotypes of some AD models. In this study we aimed to understand the relationship between Nrf2 elimination, senescence, and cognitive impairment in AD, by generating a mouse model expressing a mutant human tau transgene in an Nrf2 knockout (Nrf2KO) background. We assessed senescent cell burden and cognitive decline of P301S mice in the presence and absence of Nrf2. Lastly, we administered 4.5 month-long treatments with two senotherapeutic drugs to analyze their potential to prevent senescent cell burden and cognitive decline: the senolytic drugs Dasatinib and Quercetin (DQ) and the senomorphic drug rapamycin. Nrf2 loss accelerated the onset of hind-limb paralysis in P301S mice. At 8.5 months of age, P301S mice did not exhibit memory deficits, while P301S mice without Nrf2 were significantly impaired. However, markers of senescence were not elevated by Nrf2 ablation in any of tissues that we examined. Neither drug treatment improved cognitive performance, nor did it reduce expression of senescence markers in brains of P301S mice. Contrarily, rapamycin treatment at the doses used delayed spatial learning and led to a modest decrease in spatial memory. Taken together, our data suggests that the emergence of senescence may be causally associated with onset of cognitive decline in the P301S model, indicate that Nrf2 protects brain function in a model of AD through mechanisms that may include, but do not require the inhibition of senescence, and suggest possible limitations for DQ and rapamycin as therapies for AD.

## Introduction

Alzheimer’s Disease (AD) is an age-driven neurodegenerative disease, characterized histopathologically by accumulation of amyloid-β plaques and tau neurofibrillary tangles in the brain[1,2]. Aside from plaques and tangles, other brain changes such as brain inflammation, synaptic loss, brain atrophy, and metabolic dysregulation have been associated with AD[3]. As immunosurveillance mechanisms weaken or become overwhelmed by biological damage with age, chronic inflammation sets in and likely contributes to the development and progression of AD. Thus, understanding the underlying mechanisms aids in identifying biomarkers for early detection and therapeutics for slowing disease progression[4].

One potential source of chronic inflammation in all age-related diseases, including AD, is cellular senescence[5–7]. Cellular senescence is a cellular state characterized by an irreversible cell-cycle arrest and resistance to apoptosis that can be induced through a variety of stimuli, including telomere shortening, DNA damage, and mitotic oncogenes where cells secrete a panel of molecules deleterious to surrounding cells and tissue[8,9]. Senescent cells have been shown to accumulate in a variety of tissues with age, including the sites of chronic diseases[10]. Research has shown that senescent cells are implicated in the development of AD[11–13] and aging pathologies[14] likely due to their secretion of inflammatory molecules. Senescence is identified by a set of biomarkers characterizing different aspects of the senescence program[5,15], including: increased levels of cell cycle arrest markers (such as p16 and p21), secretion of a variety of inflammatory cytokines and other factors referred to as the senescent associated secretory phenotype (SASP), and lysosomal expansion observed as senescence associated β-galactosidase (SAβ-gal) activity. The SASP has recently emerged as a potential biomarker for early detection of AD, though cellular senescence in the brain remains largely unexplored[16]. Recently, multiple groups have demonstrated that clearance of senescent cells delays the onset of age-related pathologies (such as physical frailty, loss of endurance, and cognitive decline) in aged mice as well as in tau-based AD mouse models[6,11,17].

There are a variety of senotherapeutic treatments designed to target cellular senescence, which can be classified as either senolytic or senomorphic drugs. Senolytics, such as the combination drug dasatinib and quercetin (DQ), clear senescent cells through the induction of apoptosis. Senomorphics, such as rapamycin, prevent accumulation of senescent cells by modulating senescence markers or suppressing the secretory phenotype[12,18]. Long term DQ treatment has been shown to alleviate senescence-related phenotypes in mice and clear senescent cells from the hippocampus of aged mice, reducing cognitive impairment[10]. However, the effects of DQ on brain senescence and cognition in AD mouse models remains largely unexplored. Rapamycin has been shown to extend lifespan and healthspan in various species including mice[19–21]. mTOR attenuation has positive impacts on a variety of age-related conditions such as metabolic diseases, immune disorders, neurodegenerative diseases, and the aging process in general[20– 26]. Furthermore, rapamycin prevents senescent cell accumulation in different cell types and from multiple senescent inducers[27–30]. Studies indicate that rapamycin reduces AD-like pathology and improves cognition in the hAPP(J20)[31–33], P301S(PS19)[34], and viral vector-based P301L[35] mouse models of AD and tauopathy. Previous work in our lab showed that rapamycin can prevent, but not clear senescence in mouse embryonic fibroblasts as well as in the lung and adipose tissue of mice through the Nuclear factor-erythroid factor 2-related factor 2 (Nrf2) related pathway[27].

Nrf2, referred to as the master regulator of the redox pathway, is also a key mediator of the beneficial effects of caloric restriction, and acts as an important part of a damage response pathway and regulator of inflammation[36]. Protein levels of Nrf2 are known to decline with age, giving rise to an increased vulnerability to biological damage [37,38]. Furthermore, decreased Nrf2 levels have been observed in senescent fibroblasts and inhibition of Nrf2 promotes senescence *in vitro*[39–41]. Nrf2 knock out (Nrf2KO) mice exhibit cognitive decline, and seem to mimic the aging phenotype[42]. Similarly, breeding Nrf2KO mice with some Aβ mouse models of AD exacerbates the AD-like symptoms[43]. To date no study has assessed how the loss of Nrf2 affects brain senescence in AD mouse models.

In our study, we aimed to define how loss of Nrf2 affects cognition and brain cellular senescence of transgenic MAPT^P301S^PS19 (P301S) mice, which expresses high levels of P301S mutant human tau specifically in neurons. This model exhibits neurofibrillary tangle (NFT) deposition, neurodegeneration and impaired cognitive function[6] as well as severe hind-limb paralysis and an inability to feed, resulting in death at an early age[44]. We also aimed to determine the potential of high-dose rapamycin and DQ to clear senescence and improve cognition in P301S mice in the presence and absence of Nrf2. To this end we bred Nrf2KO mice with transgenic P301S mice to generate non-transgenic Nrf2 wildtype (WT), Nrf2KO, transgenic Nrf2 wildtype (TgWT), and transgenic Nrf2KO (TgKO) mice, and treated transgenic mice with either DQ, high-dose rapamycin, or vehicle from 4 to 8.5 months of age. All WT mice were treated with vehicle. Cognition was assessed through MWM testing at 8.5 months. Then hippocampus, prefrontal cortex, whole brain, lung, and adipose tissue, along with astrocytes isolated from brain, were collected for analysis of senescent markers. We found that loss of Nrf2 greatly accelerated the development of hind-limb ataxia and paralysis in P301S mice from 13 to 10 months of age. TgKO mice showed long-term memory impairment at 8.5 months of age, while TgWT mice did not. However, loss of Nrf2 did not elevate levels of senescence in any tissue of P301S mice. Further, both high-dose rapamycin and DQ failed to improve cognitive performance or reduce senescent cell burden in the presence or absence of Nrf2. High-dose rapamycin instead seemed to impair cognition and potentially elevate levels of SAβ-gal activity.

## Materials and Methods

### Mouse Strains

This study received prior approval from Oregon State University Institutional Animal Care and Use Committee. MAPT^P301S^PS19 (P301S) mice (JAX stock #008169)[44] were purchased from The Jackson Laboratory and bred to C57BL/6J for six generations. Nrf2KO mice (ICR background) were obtained from Dr. Masayuki and Yamamoto Kohoku, University of Japan, and then were backcrossed 10 times into the C57BL6/J genetic background. P301S mice were bred with Nrf2KO over multiple generations to generate non-transgenic Nrf2 wildtype (WT), transgenic Nrf2 wildtype (TgWT), and transgenic Nrf2KO (TgKO) mice.

### Animal Treatment

For both the DQ and rapamycin experiments, mice were divided into five groups, each containing between 11 and 15 mice per sex. For the senolytic experiment all WT mice received vehicle treatment, while TgWT and TgKO mice were randomly divided into drug and vehicle treatment groups. Treatment began at 4.5 months of age, continuing for five days a week, every other week until 8.5 months of age. Mice received Dasatinib and Quercetin by oral gavage at a dose of 5 and 50 mg kg^-1^ respectively diluted into vehicle or with the same volume of vehicle as controls[45–48]. For the rapamycin experiment all WT mice received vehicle treatment, while TgWT and TgKO mice were randomly divided into drug and vehicle treatment groups.

Treatment began at 4.5 months of age, continuing every other day until 8.5 months of age. Mice received rapamycin by I.P. injection at a dose of 8 mg kg^-1^ diluted into vehicle or with the same volume of vehicle as controls[49]. This dose was used as intervention for short-lived models of dilated cardiomyopathy, muscular dystrophy and Leigh Syndrome[26], a severe mitochondrial disease[50] and in short-intervention lifespan studies[49], and is approximately 4-fold higher than that used in lifespan studies[20–22] and in studies of rapamycin in mouse models of AD[25,31–33,51,52]. Mice were euthanized 48 h after the last dose of rapamycin, DQ, or vehicle and a thorough necropsy was performed. Tissues were snap frozen in liquid nitrogen, then stored at -80 °C for subsequent molecular work.

To determine onset of hind-limb ataxia, cohorts of WT (n=10), TgWT (n=11), TgKO (n=11), and Nrf2KO (n=4) mice were observed twice a week for evidence of weight loss and hind limb clamping and the age of phenotypic onset was recorded. The data were plotted as a survival curve using GraphPad Prism software and analyzed by log-rank test (Mantel-Cox test) comparing the TgWT and TgKO groups.

### Morris Water Maze

Male and female P301S(PS19) and P301S X Nrf2KO mice at 8.5 months of age were tested in the Morris Water Maze (MWM) task, including acclimation (2 days), spatial memory testing, including 16 reference hidden trials or place trials, and a naive probe (ran without knowledge of platform location) and other probes at the beginning of every other day (4 days); reversal training (1 day) and visible (control) trials (1 day) as previously described[53]. Performance on the spatial and cued version of the MWM task was analyzed using Any-maze software. Performance was measured as the corrected integrated path length (CIPL)[54,55] for spatial learning and reversal trials. CIPL, rather than latency or distance swam, is reported to avoid the variability in results introduced by differences in swimming speed and release locations[53]. Mean distance from the former platform location was used as a measure of spatial memory on probe trials. Performing probes at multiple points throughout training allows us to see a progression of memory formation, although it can negatively impact learning. Animals may not show any differences in probe trials after training, but may show it early on[53]. N = 11-14 mice per group. Analyses were performed with repeated measures ANOVA (Fishers LSD post-hoc tests were performed as indicated when main effects of place trial or treatment/genotype were observed) using Prism 6 (GraphPad, San Diego, CA, USA).

### RT-qPCR analysis

mRNA was extracted using the RNeasy kit (Qiagen, Valencia, CA, USA). RNA was reverse-transcribed to cDNA using SuperScript® IV FirstStrand Synthesis SuperMix following the manufacturer’s instructions (Invitrogen). Target mRNA levels were measured by qPCR and normalized to beta actin (actb). Amounts of specific mRNA were quantified by determining the point at which fluorescence accumulation entered the exponential phase (Ct), and the Ct ratio of the target gene to actb was calculated for each sample. Ct ratios were further normalized to average WT expression. Information for qPCR primers is listed in Supplementary Table 3.1.

### Western Blotting

Brain tissue was homogenized in RIPA buffer supplemented with protease and protein phosphatase inhibitors (Calbiochem, La Jolla, CA) and subjected to SDS-PAGE followed by transferring to PVDF membranes (Millipore, Billerica, MA, USA). Membranes were incubated with antibodies specific for: phosphor-Tau (Ser 199, Ser 202), Tau monoclonal antibody (Tau-5) (Thermo Fisher Scientific, Waltham, MA), and Actin (MP Biomedicals, Solon, OH). The intensity of the bands was quantified by densitometry using Imagelab software (Bio-rad, Hercules, CA).

### SAβ-gal staining

SA-βgal staining was performed on coronal cryosections of hemibrains using the SPiDER-βgal kit (Dojindo Molecular Technologies, Rockville, MD, USA). Briefly, brain sections in groups of 3 were fixed onto slides with 4% paraformaldehyde, incubated with 80 µl of SPiDER-βgal working solution at 37°C for 2.5 hours, washed 3 times with PBS, then incubated with 4′,6-diamidino-2-phenylindole (DAPI), a fluorescent stain that binds tightly to adenine-thymine rich regions of DNA at 4°C overnight, and imaged (excitation: 488 nm; emission: 500-600 nm) using a Keyence fluorescence microscope.

### Statistical analysis

QPCR and western blot data were analyzed using Prism 6 (GraphPad, San Diego, CA, USA). Significance of differences between experimental group means were defined using one-way ANOVA. Fisher’s LSD tests were performed to ascertain differences between genotype/treatment means, and P < 0.05 was considered significant and reported as indicated in the legends to the figures.

## Results

### Nrf2 loss accelerated development of hind-limb paralysis in P301S mice

P301S mice express high levels of mutant human tau (hTau), roughly 5-fold higher than endogenous mouse tau, resulting in rapidly lethal, early onset disease that is characterized by a severe hind-limb dystonia, paralysis, an inability to feed, and death at 9 months of age[44]. We observed a cohort of 10-12 animals, WT, TgWT, TgKO, and Nrf2KO mice over a period of 15 months after birth to generate a survival curve based on the onset of hind-limb paralysis (**Figure 1**). Over the 15-month period, TgKO mice developed hind-limb atrophy between 8.5 and 11 months of age, with 50% animals euthanized by 10 months of age. This was compared to TgWT mice that developed hind-limb atrophy between 10.5 and 15 months of age, with 50% of the group euthanized by 13 months of age (**Figure 1**). No Nrf2KO mice and only one WT mouse were euthanized over the 15-month period. Only 4 Nrf2KO mice were observed for this study, however mice of this genotype do not display hind-limb dysfunction at least until 20 months of age in other studies [56]. These results indicate that hind-limb ataxia was delayed in TgWT mice compared to previously reported studies, and that loss of Nrf2 significantly accelerated the hind limb atrophy in the TgKO mice compared to the TgWT group.

**Figure 1.**
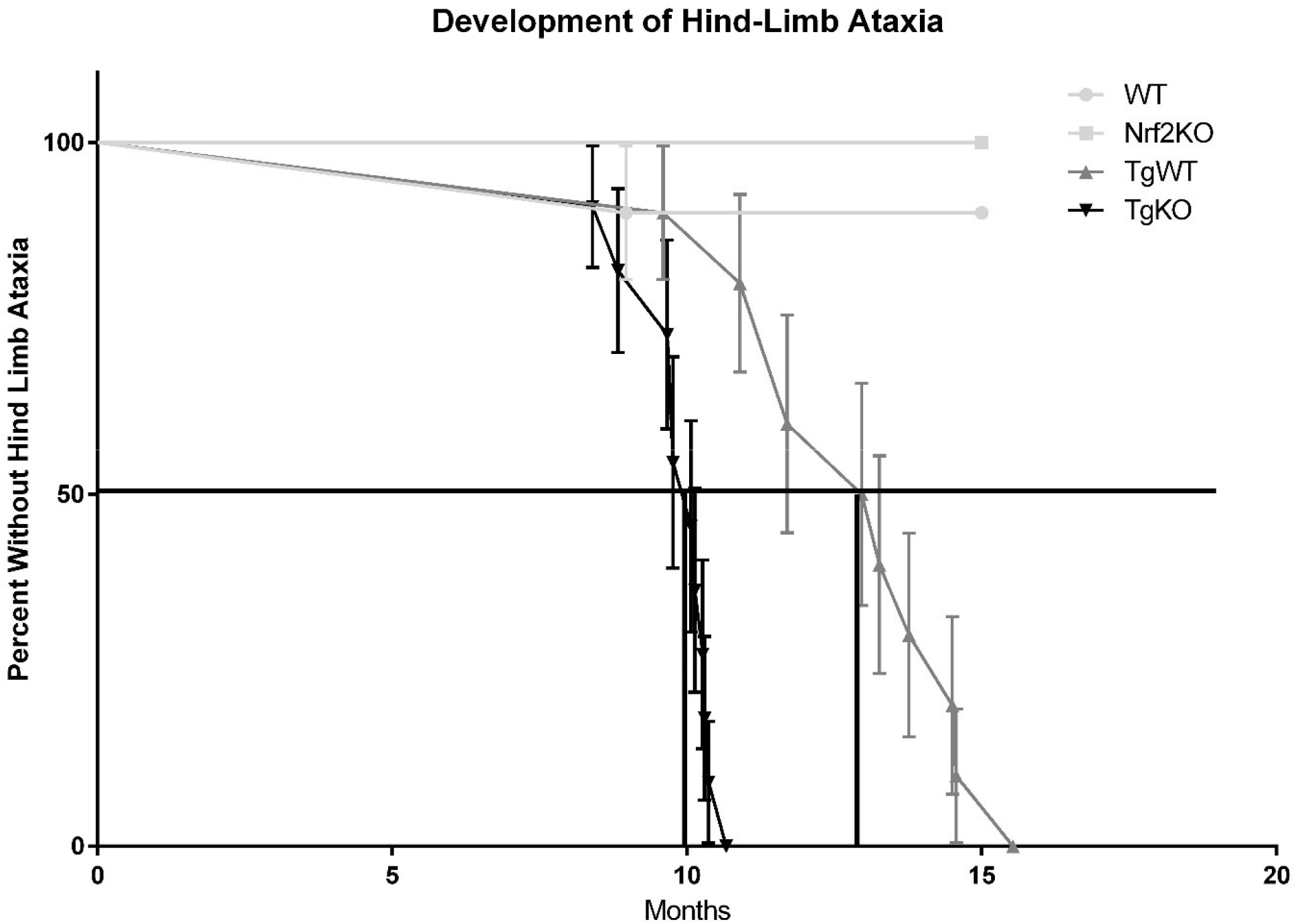
Loss of Nrf2 accelerated the development of hind limb ataxia in P301S mice. Hind-limb clamping was observed and recorded weekly up to 8 months of age, and then monitored every other day up to 15 months of age. The light gray line with squares represents Nrf2KO mice, the light gray line with circles represents WT mice, the gray line with triangles represents TgWT mice, and the black line with triangles represents TgKO mice. Each point represents death or the time at which hind-limb phenotype appeared, requiring mice to be euthanized. TgKO mice developed hind limb ataxia at 10 months of age, which was significantly earlier than TgWT mice at 13 months of age (Log rank Mantel Cox test, p<0.0001 for difference between TgWT and TgKO curves). Nrf2KO and WT mice largely did not experience the hind-limb phenotype, with 100 and 90% surviving past 15 months of age respectively. TgKO n=11, TgWT n=10, WT n=10, Nrf2KO n=4

### Nrf2 loss did not increase pTau levels in P301S brain

Elevated levels of tau and phosphorylated tau (pTau) in the spinal cord of P301S mice results in severe hind-limb ataxia[44], which is accelerated by loss of Nrf2. We analyzed the effect of Nrf2 ablation on tau and pTau levels in brains of P301S mice to determine whether acceleration of hindlimb ataxia correlates with increased pTau burden. Western blot analysis of P301S mouse brains showed a roughly 12-fold increase in tau levels and 4-fold increase of pTau levels compared to WT controls. However, loss of Nrf2 did not increase either tau or pTau in TgKO mice compared to the TgWT group (**Figure 2**).

**Figure 2.**
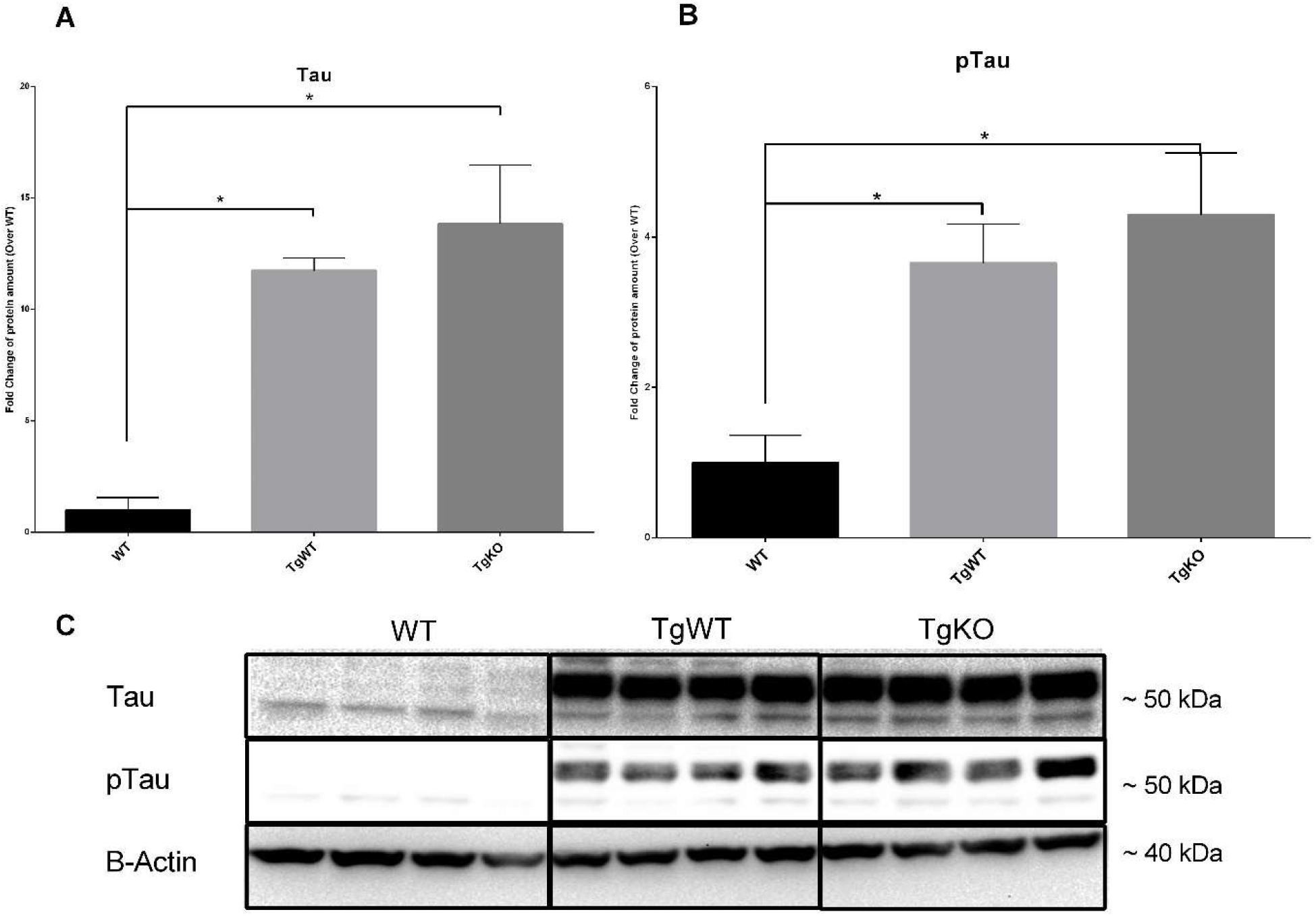
Nrf2 ablation did not elevate brain levels of tau or ptau. Western blot analysis of tau (Figure A) and ptau (Figure B) protein expression in mouse brains at 8.5 months of age using anti-Tau monoclonal antibody (Tau-5) and anti-phosphoTau (Ser 199, Ser 202) antibody, normalized first to β-actin and then WT expression. Representative western blot, Tau and pTau appear ∼ 50-55 kDA, and β-actin appears at ∼ 37-40 kDA (Figure C). *p<0.05 for differences between genotypes, one way ANOVA with multiple comparisons, data = mean ± SEM)

### Loss of Nrf2 accelerated long term memory decline of P301S mice

Our next goal was to determine if loss of Nrf2 worsens the cognitive decline observed in the P301S model. While it is common to measure senescence and cognition by MWM at 6 months of age in this model, our previous studies (Dorigatti et al 2021) showed that in our hands, P301S did not exhibit senescence or cognitive decline at 6-7 months of age but developed senescence at 10 months of age[15]. Given that TgKO mice begin developing hind limb ataxia at 8.5 months of age, we judged 8.5 months to be the best age to measure senescence and cognition.

To assess spatial learning and memory and cognitive flexibility, we used the MWM on WT, TgWT, and TgKO mice at 8.5 months of age as described[53]. We found significant main effects of genotype and place trial in the MWM tasks (**Table 1**). There was a main effect of place trial in the spatial learning task for both male (p<0.0001) and female mice (p=0.0138). With data collapsed across genotype, both sexes showed lower CIPL measurements in the last trials as compared to the first (p<0.0001).

There was a significant main effect of genotype on CIPL scores for spatial learning in female mice (p<0.001, **Table 1**): TgKO mice had significantly higher CIPL scores on the spatial learning trials than WT mice (p=0.0484), while TgWT mice did not differ from WT (p=0.3485; **Figure 3 a-b**). There was no significant main effect of genotype in the spatial learning task for males (p=0.1506; **Table 1**; **Figure 3 c-d**).

**Figure 3.**
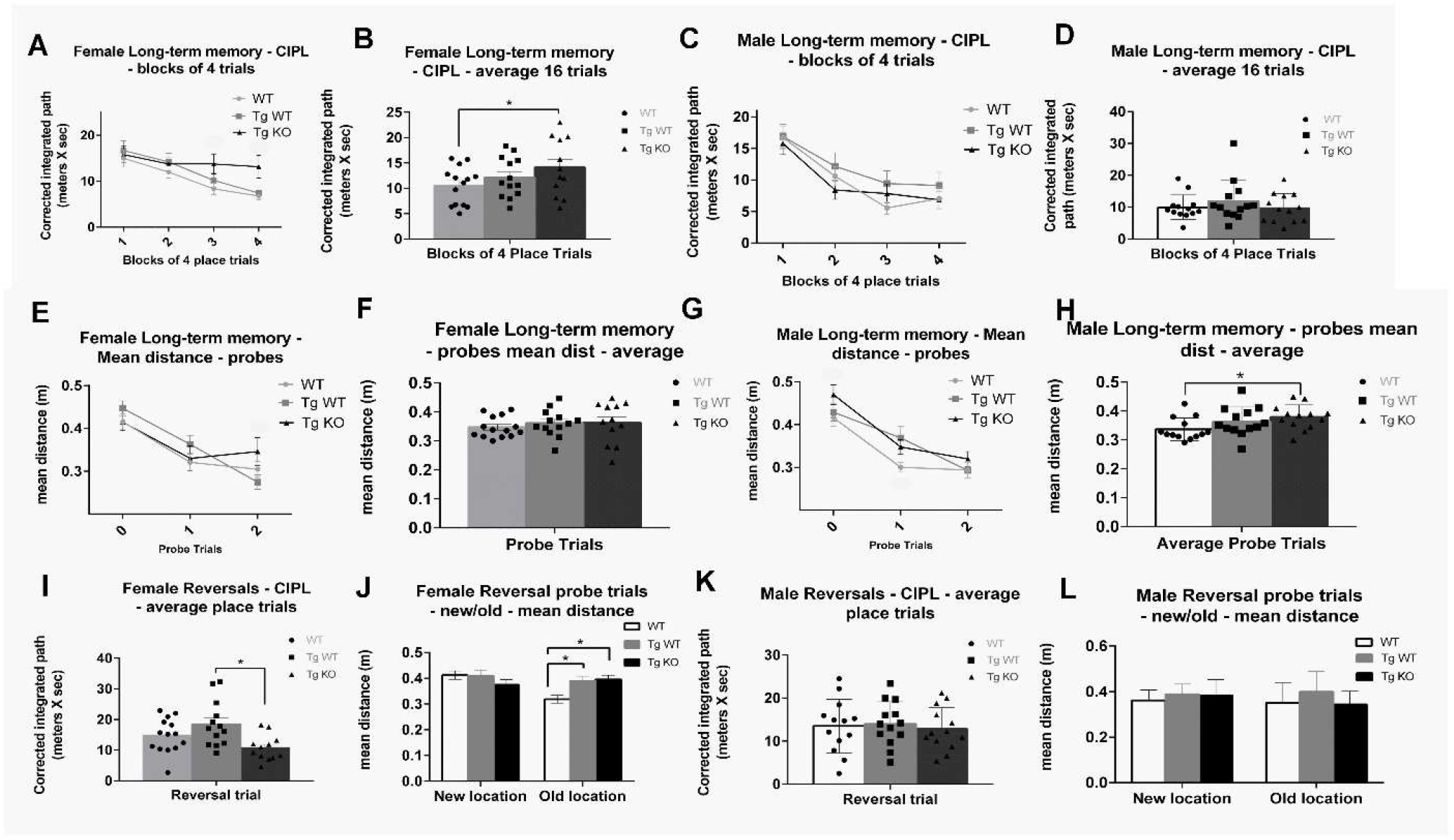
Nrf2 elimination accelerated long term memory decline in P301S mice. Analysis of MWM data for spatial learning, spatial memory and cognitive flexibility for female and male WT (Light gray lines and bars with circles), TgWT (Gray lines and bars with squares), and TgKO (Black lines and bars with triangles) mice at 8.5 months of age. Spatial learning ability (Figures A-D) of females and males reported as corrected integrated pathlength (CIPL), with higher values indicating worse performance. Memory of platform location (Figures E-H) measured by mean distance from former platform location. Cognitive flexibility (Figures I-L) determined by reversal place trials and new/old platform location probes. (n=11-14 per group, *p<0.05 for differences between genotypes defined by Fishers LSD post-hoc test on a significant effect of genotype, ANOVA, data = mean ± SEM)

**Table 1.**
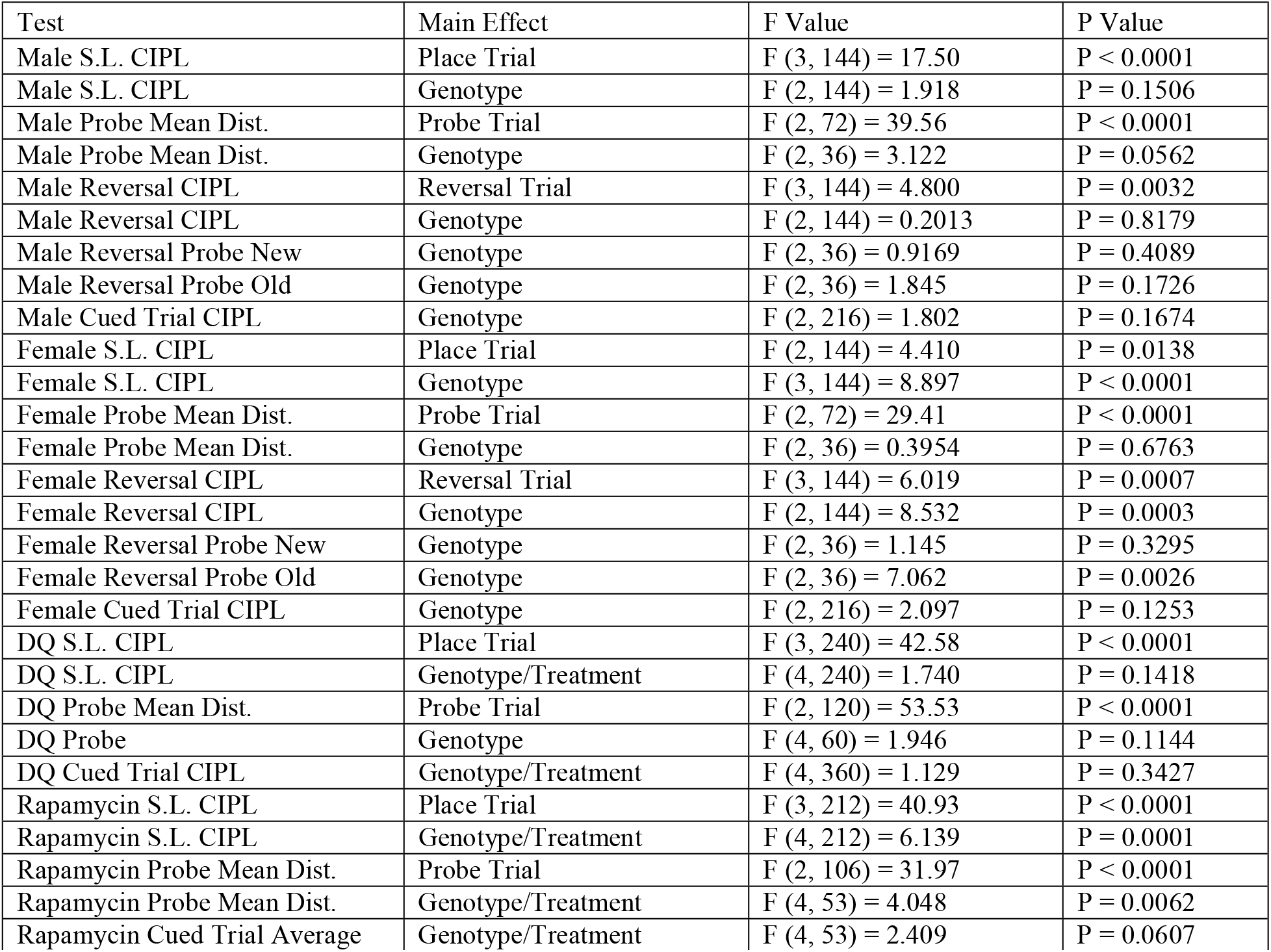
Two Way ANOVA Main Effects for MWM. F statistics and P values for main effect of MWM testing. Tests include Spatial learning trials (S.L.) by corrected integrated path length (CIPL), long-term memory Probes by mean distance, reversal trials by CIPL, and reversal probes by mean distance of new and old locations. Fishers LSD post-hoc tests were performed as indicated when main effects of place trial or treatment/genotype were observed

| Test | Main Effect | F Value | P Value |
| --- | --- | --- | --- |
| Male S.L. CIPL | Place Trial | F (3, 144) = 17.50 | P < 0.0001 |
| Male S.L. CIPL | Genotype | F (2, 144) = 1.918 | P = 0.1506 |
| Male Probe Mean Dist. | Probe Trial | F (2, 72) = 39.56 | P < 0.0001 |
| Male Probe Mean Dist. | Genotype | F (2, 36) = 3.122 | P = 0.0562 |
| Male Reversal CIPL | Reversal Trial | F (3, 144) = 4.800 | P = 0.0032 |
| Male Reversal CIPL | Genotype | F (2, 144) = 0.2013 | P = 0.8179 |
| Male Reversal Probe New | Genotype | F (2, 36) = 0.9169 | P = 0.4089 |
| Male Reversal Probe Old | Genotype | F (2, 36) = 1.845 | P = 0.1726 |
| Male Cued Trial CIPL | Genotype | F (2, 216) = 1.802 | P = 0.1674 |
| Female S.L. CIPL | Place Trial | F (2, 144) = 4.410 | P = 0.0138 |
| Female S.L. CIPL | Genotype | F (3, 144) = 8.897 | P < 0.0001 |
| Female Probe Mean Dist. | Probe Trial | F (2, 72) = 29.41 | P < 0.0001 |
| Female Probe Mean Dist. | Genotype | F (2, 36) = 0.3954 | P = 0.6763 |
| Female Reversal CIPL | Reversal Trial | F (3, 144) = 6.019 | P = 0.0007 |
| Female Reversal CIPL | Genotype | F (2, 144) = 8.532 | P = 0.0003 |
| Female Reversal Probe New | Genotype | F (2, 36) = 1.145 | P = 0.3295 |
| Female Reversal Probe Old | Genotype | F (2, 36) = 7.062 | P = 0.0026 |
| Female Cued Trial CIPL | Genotype | F (2, 216) = 2.097 | P = 0.1253 |
| DQ S.L. CIPL | Place Trial | F (3, 240) = 42.58 | P < 0.0001 |
| DQ S.L. CIPL | Genotype/Treatment | F (4, 240) = 1.740 | P = 0.1418 |
| DQ Probe Mean Dist. | Probe Trial | F (2, 120) = 53.53 | P < 0.0001 |
| DQ Probe | Genotype | F (4, 60) = 1.946 | P = 0.1144 |
| DQ Cued Trial CIPL | Genotype/Treatment | F (4, 360) = 1.129 | P = 0.3427 |
| Rapamycin S.L. CIPL | Place Trial | F (3, 212) = 40.93 | P < 0.0001 |
| Rapamycin S.L. CIPL | Genotype/Treatment | F (4, 212) = 6.139 | P = 0.0001 |
| Rapamycin Probe Mean Dist. | Probe Trial | F (2, 106) = 31.97 | P < 0.0001 |
| Rapamycin Probe Mean Dist. | Genotype/Treatment | F (4, 53) = 4.048 | P = 0.0062 |
| Rapamycin Cued Trial Average | Genotype/Treatment | F (4, 53) = 2.409 | P = 0.0607 |

Probe trials were run at the beginning, middle, and end of the spatial learning process in which the platform was removed to test memory of the platform location. For female mice there was no significant main effect of genotype in the probe trials (p=0.6763; **Table 1**; **Figure 3 e-f**). Males showed a trend for main effect of genotype on mean distance scores for probe trials (p=0.0562; **Table 1**): TgKO mice had significantly higher mean distance scores on probe trials than WT mice (p=0.0184), while TgWT mice did not differ from WT (p=0.1262; **Figure 3 g-h**).

To test cognitive flexibility, the platform location was moved, and mice were given four learning trials to learn the new platform location followed by a probe for the new and old platform locations. Amongst males, there was no significant main effect of genotype for reversal trials or the probe for new or old platform location (p=0.8179-0.1726; **Table 1**; **Figure 3 i-j**). There was a significant main effect of genotype on CIPL scores for female mice on reversal trials (p=0.0003; **Table 1**): Neither TgWT nor TgKO mice had significantly different CIPL scores than WT (p=0.1154-0.0896; **Figure 3 k-l**).

Our data showed no main effect of genotype on CIPL scores for cued trials of either sex, indicating no visual deficiency (p=0.1674-0.1253; **Table 1**; **supplementary figure 2**).

### Loss of Nrf2 did not elevate levels of hippocampal senescence

In previous studies, the elimination of Nrf2 resulted in increased cellular senescence *in vitro* and *in vivo*[27]. To define the effect of Nrf2 loss on the accumulation of senescent cells in the hippocampus of P301S mice, we performed QPCR analysis on cell cycle arrest markers and common SASP factors from flash frozen hippocampus tissue collected from 8.5 month-old WT, TgWT, and TgKO mice, as well as SAβ-gal staining on hippocampus-containing brain cryo-sections from the same groups.

QPCR analysis of hippocampal tissue showed significant main effects of genotype for expression of SASP factors (F (2, 205) = 9.090; p=0.0002) and cell cycle arrest markers (F (2, 65) = 7.283; p=0.0014): TgWT and TgKO mice had significantly elevated levels of IL1-α, IL1-β, and p16 (p=0.0288-<0.0001) compared to WT mice. TgWT and TgKO mice also showed trends to elevations in ICAM1 and p21 expression (p=0.0669 and p=0.0611 respectively). However, TgKO expression of all SASP factors and cell cycle arrest markers was similar to TgWT mice (p=0.998-0.7785; **Figure 4 a-b**). This analysis was repeated in isolated brain astrocytes as well as in whole prefrontal cortex and in adipose tissue, yielding no significant increases of senescence markers in P301S mice regardless of Nrf2 status (**Supplementary Figure 1 a-j**).

**Figure 4.**
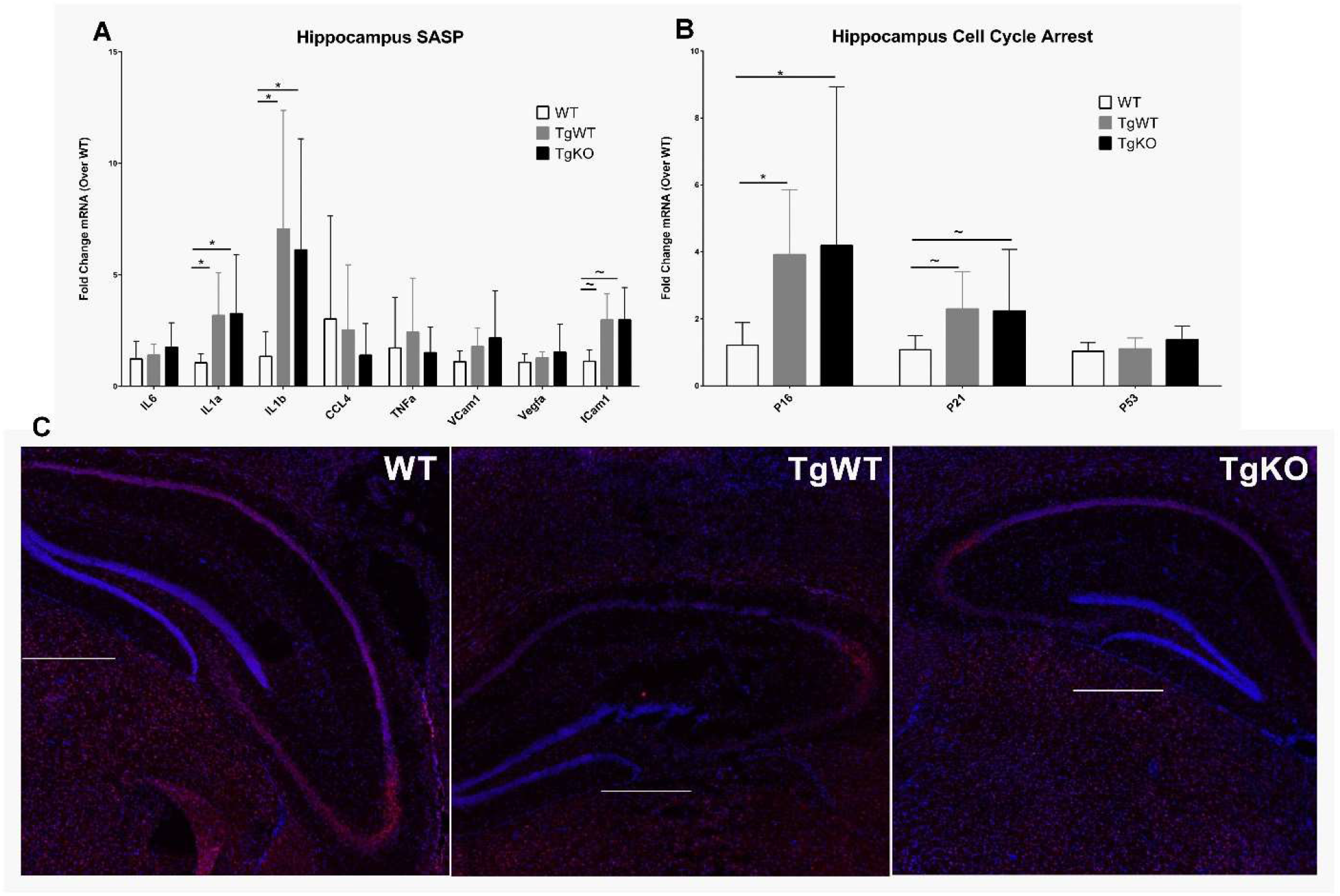
Nrf2 loss did not exacerbate senescent cell burden in the hippocampus of P301S mice. Senescent cell burden was determined by analysis of 3 biomarkers: SASP, cell cycle arrest, and SAβ-gal activity. qPCR analysis of SASP (A) and cell cycle arrest markers (B) mRNA expression in hippocampus tissue of WT (White bars), TgWT (Gray bars), and TgKO (Black bars) mice at 8.5 months of age. Analysis of Saβ-gal activity (C), indicated by red fluorescence (total cell number, blue fluorescence), in brain cryosections from WT, TgWT or TgKO mice. (Scale bar = 500 µm. *p<0.05, ∼p<0.07 for differences between genotypes, one way ANOVA with multiple comparisons using Fishers LSD post-hoc test, n=8-11 per group at 8.5 months of age, data = means ± SEM)

SAβ-gal staining showed no increase in β-gal activity in hippocampi of TgWT or TgKO mice compared to WT mice. Similarly, β-gal activity was similar for TgWT mice compared to TgKO (**Figure 4 c**).

Due to the lack of senescent cell accumulation, we searched for evidence of brain inflammation through expression of GFAP to indicate astrocyte activation. QPCR analysis showed a genotype effect of GFAP expression for the hippocampus (F (4, 16) = 3.567; p=0.0291): TgWT (p=0.0113) and TgKO (p=0.0056) mice expressed higher levels of GFAP than WT mice. However, TgKO mice showed similar levels of GFAP to TgWT (p=0.7537; **Supplementary Figure 1 k-l**).

### Effect of senotherapeutic treatment on cognition

We aimed to determine the effects of the senotherapeutic treatments, DQ and rapamycin, on cognition of TgWT and TgKO mice by MWM. For both therapeutics, either drug or vehicle treatment began at 4.5 months of age until completion of MWM testing at 8.5 months of age. We found no significant main effects of genotype/treatment for spatial learning trials or long-term memory probes of DQ treated mice (p=0.1418-0.1144; **Table 1**; **Figure 5 a-d**). For rapamycin treatment, we found a main effect of genotype/treatment for CIPL scores of spatial learning trials (p=0.0001; **Table 1**): Rapamycin treated TgWT mice had significantly higher CIPL scores than vehicle treated WT (p=0.0065) and TgWT mice (p=0.0442; **Figure 6 a-b**). There was also a significant main effect of genotype/treatment for mean distance scores on spatial memory probes for rapamycin treated mice (p<0.0001; **Table 1**): Rapamycin treated TgWT mice had significantly higher mean distance scores than vehicle treated WT (p=0.0002) and TgWT mice (p=0.0366). Rapamycin treated TgKO mice had significantly higher mean distance scores than vehicle treated WT mice (p=0.0414), while vehicle treated TgKO mice did not differ from WT (p=0.1563; **Figure 6 c-d**).

**Figure 5.**
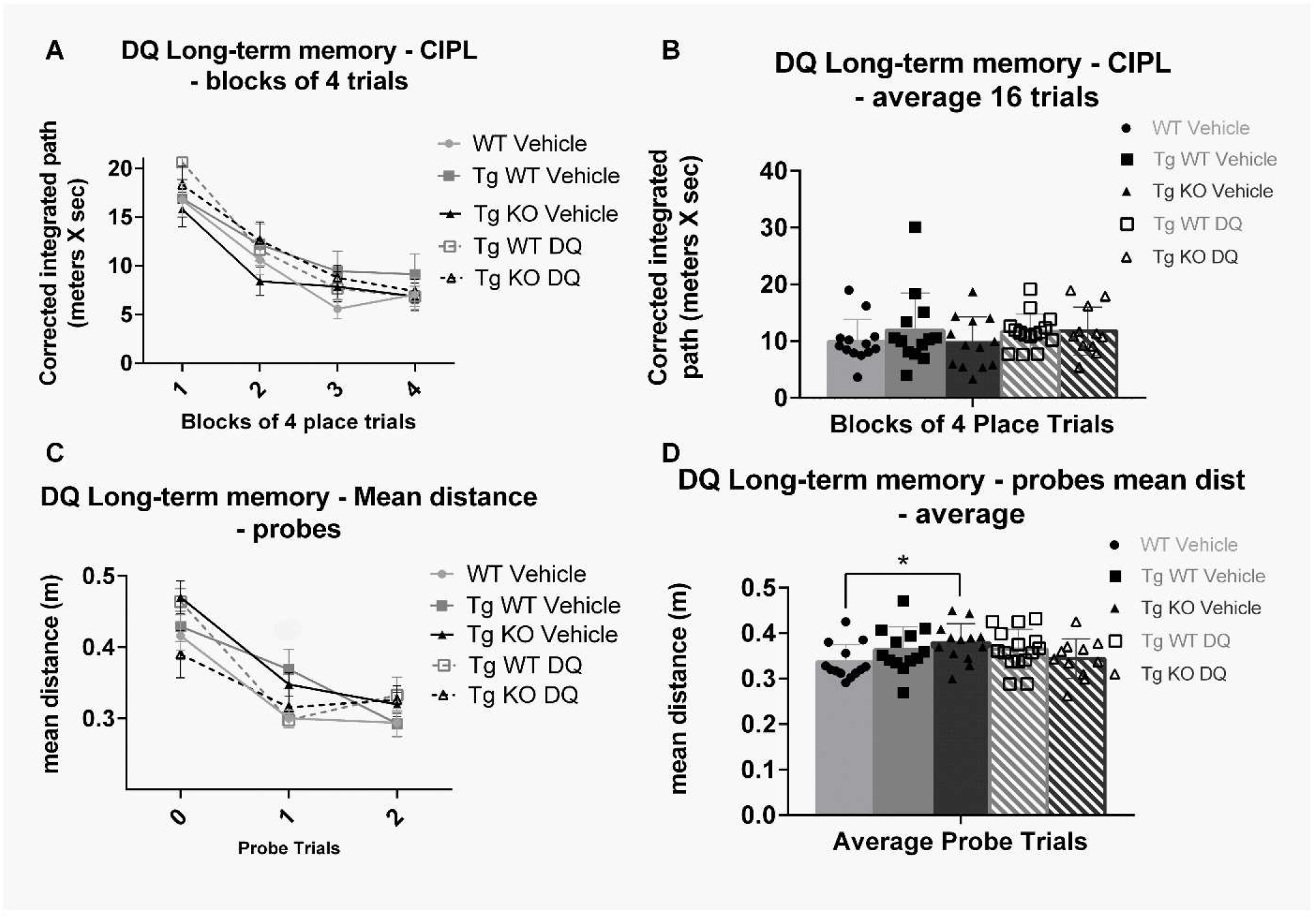
DQ treatment did not improve cognitive performance in P301S mice. MWM analysis of long-term memory of DQ treated mice (WT vehicle =light gray lines and bars with circles, TgWT vehicle=gray lines and bars with squares, TgKO vehicle=black lines and bars with triangles, TgWT DQ=striped gray with squares, TgKO DQ=striped black with triangles). Spatial learning trials of DQ and vehicle treated mice shown in blocks of four (A) and average of 16 trials (B). Long term memory probes of DQ and vehicle treated mice shown in individual probes (C) and average of 3 probes (FD). (n=11-14 per group, *p<0.05 for differences between genotypes or treatment defined by Fishers LSD post-hoc test on a significant effect of genotype, ANOVA, data = mean ± SEM)

**Figure 6.**
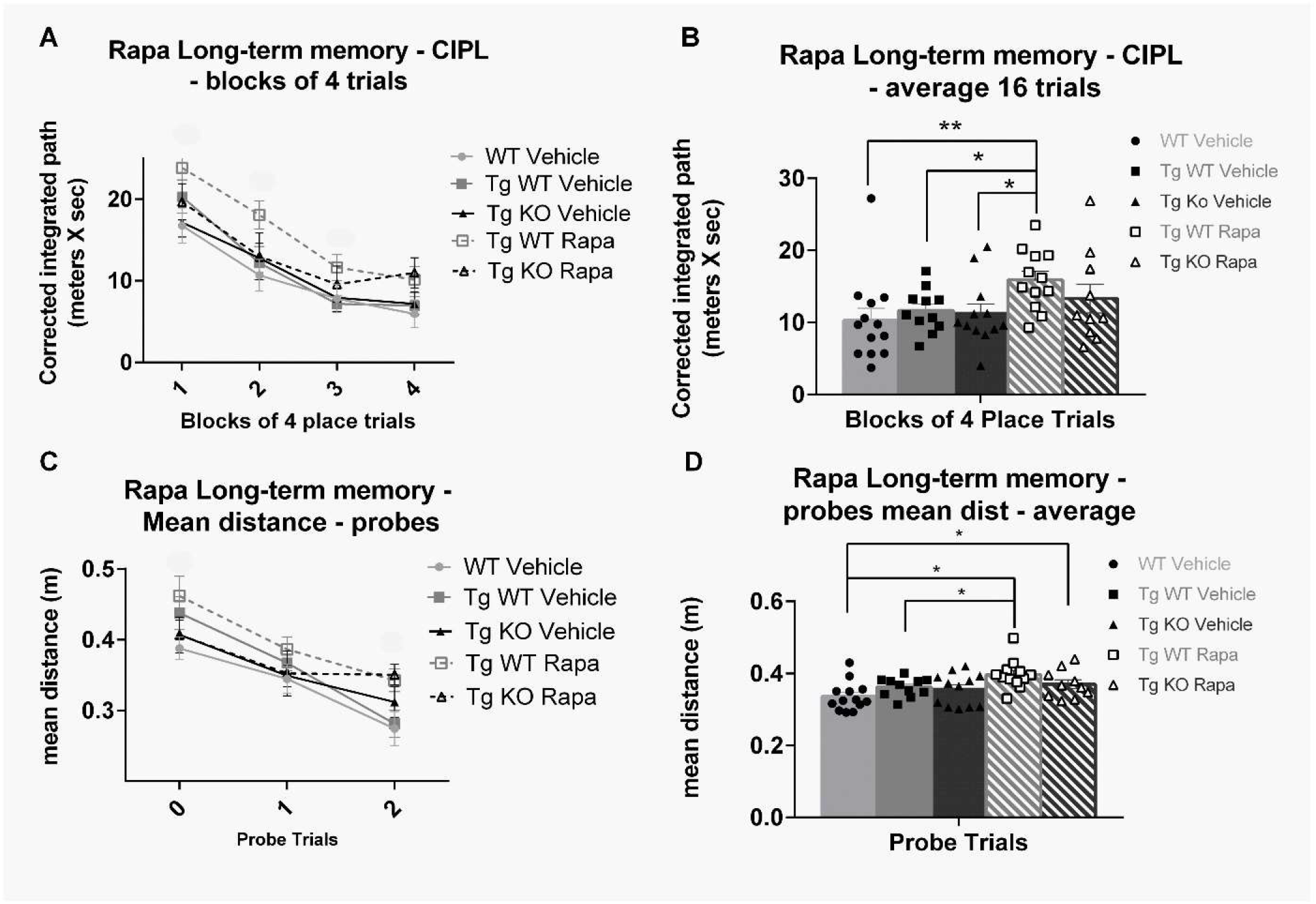
Rapamycin treatment impaired cognition in P301S mice. MWM analysis of spatial learning and memory of rapamycin treated mice (WT vehicle =light gray lines and bars with circles, TgWT vehicle=gray lines and bars with squares, TgKO vehicle=black lines and bars with triangles, TgWT rapamycin=striped gray with squares, TgKO rapamycin=striped black with triangles). Spatial learning trials of rapamycin and vehicle treated mice shown in blocks of four (A) and average of 16 trials (B). Spatial memory probes of rapamycin and vehicle treated mice shown in individual probes (C) and average of 3 probes (D). (n=11-14 per group, *p<0.05 for differences between genotypes or treatment defined by Fishers LSD post-hoc test on a significant effect of genotype, ANOVA, data = mean ± SEM).

### Effect of senotherapeutic treatment on brain senescence

Hippocampal QPCR analysis of P301S mice treated with DQ showed a significant main effect of genotype/treatment for SASP (F (4, 338) = 7.033; p<0.001) and cell cycle arrest markers (F (4, 104) = 4.087; p=0.0041): DQ treated TgWT and TgKO mice expressed similar levels of SASP factors and cell cycle arrest markers compared to respective vehicle treated mice (p=0.8052-0.0953; **Figure 7 a-b**), except for IL1β, which was reduced in DQ treated TgKO mice compared to vehicle treated TgKO mice (p<0.0001). SAβ-gal staining on brain cryosections of DQ treated P301S mice showed no change in β-gal activity compared to vehicle treated transgenic mice, independent of Nrf2 status (**Figure 7 c**).

**Figure 7.**
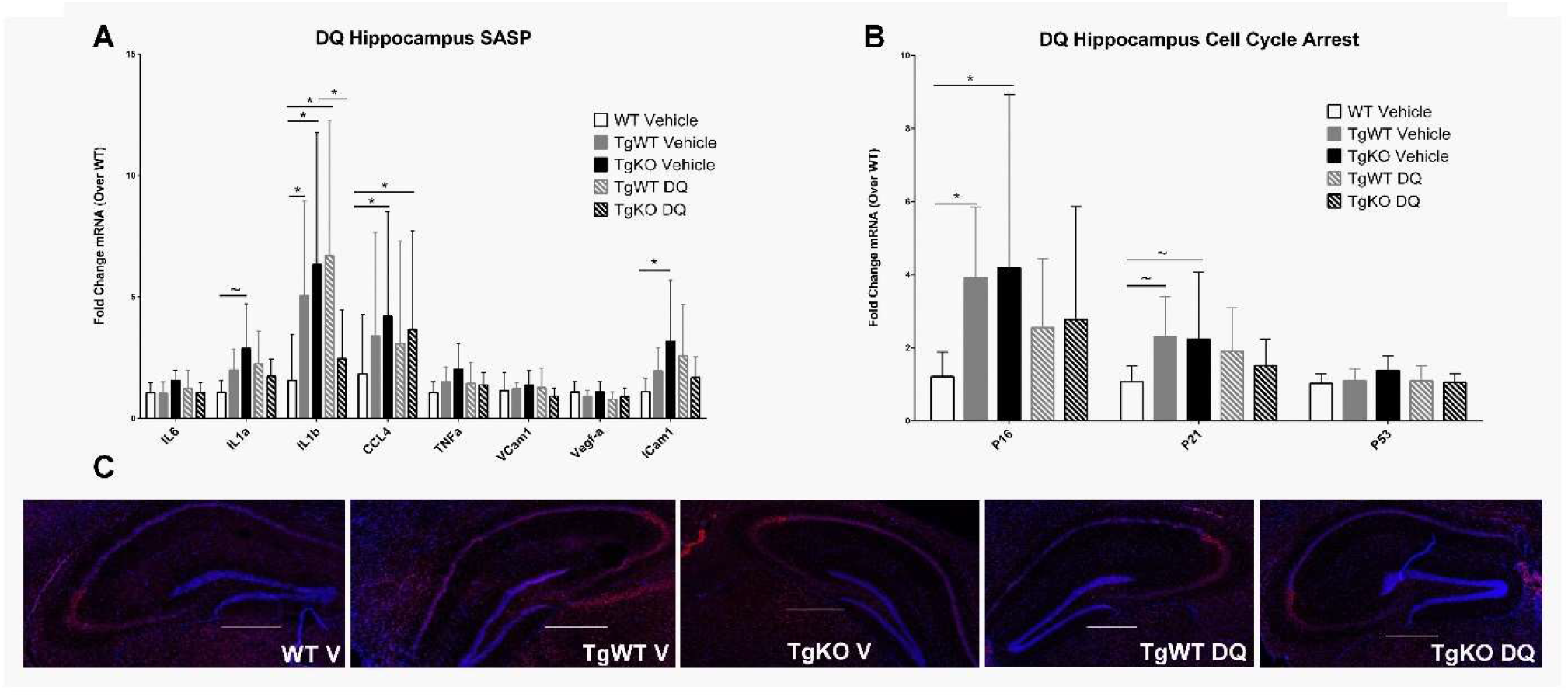
DQ treatment did not reduce markers of senescence in the hippocampus of P301S mice. Senescent burden was determined by analysis of 3 biomarkers: SASP, cell cycle arrest, and SAβ-gal activity. qPCR analysis of SASP (Figure A) and cell cycle arrest markers (Figure B) mRNA expression in hippocampus tissue of Wt Vehicle (white bars), TgWT Vehicle (Gray bars), TgKO Vehicle (Black bars), TgWT DQ (Striped gray bars), and TgKO DQ (Striped black bars) mice at 8.5 months of age. Analysis of Saβ-gal activity (Figure C), indicated by red fluorescence (total cell number, indicated by blue fluorescence), in brain cryo-sections from WT, TgWT or TgKO mice. (Scale bar = 500 µm. *p<0.05, ∼p<0.06 for differences between genotype defined by Fishers LSD post-hoc test on a significant effect of genotype, one way ANOVA with multiple comparisons, n=8-11 per group at 8.5 months of age, data = means ± SEM)

Hippocampal QPCR of transgenic mice treated with rapamycin showed a significant main effect of expression for SASP (F (4, 104) = 4.087; p=0.0009), and a near significant main effect for cell cycle arrest markers (F (4, 104) = 2.248; p=0.068): SASP and cell cycle arrest marker expression of rapamycin treated TgKO and TgWT mice were similar to respective vehicle treated mice (p=0.9826-0.3244; **Figure 8 a-b**), except for IL1β which was reduced in rapamycin treated TgKO mice compared to vehicle treated TgKO mice (p=0.0313). SAβ-gal staining on brain cryosections of rapamycin treated transgenic mice showed an increase in β-gal activity compared to vehicle treated transgenic mice in both the presence and absence of Nrf2 (**Figure 8 c**). Because mTOR is a repressor of the lysosomal pathway[57,58], decreased mTOR activity resulting from rapamycin treatment likely contributes to lysosomal expansion, observed as increased SAβ-gal activity. Further analysis of isolated astrocytes, prefrontal cortex, and adipose tissue showed no significant differences in senescent marker expression between drug and vehicle treated TgWT or TgKO mice (**Supplementary Figure 1 a-j**).

**Figure 8.**
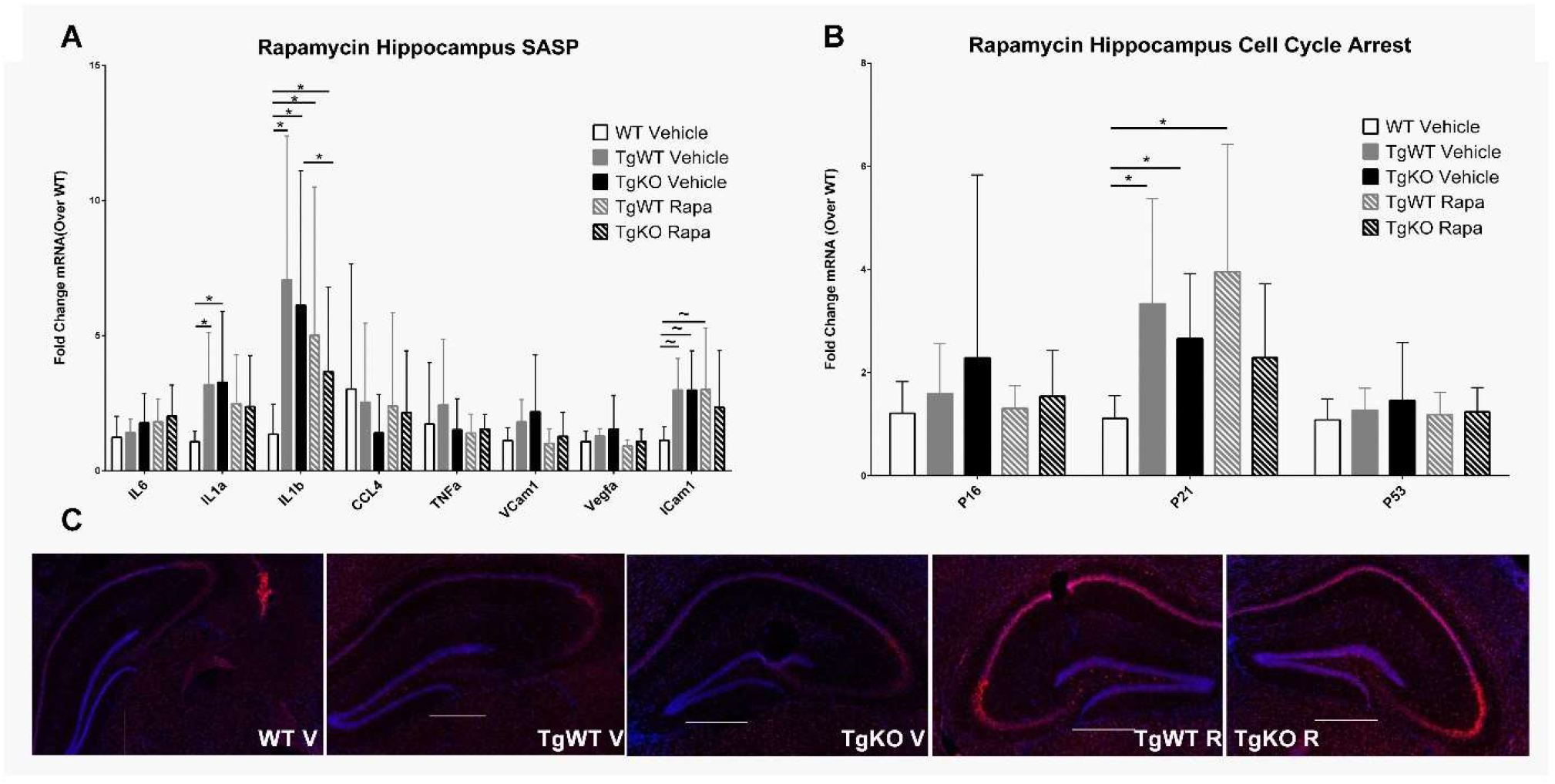
Rapamycin treatment did not reduce markers of senescence in the hippocampus of P301S mice. Senescent burden was determined by analysis of 3 biomarkers: SASP, cell cycle arrest, and SAβ-gal activity. qPCR analysis of SASP (Figure A) and cell cycle arrest markers (Figure B) mRNA expression in hippocampus tissue of Wt Vehicle (White bars), TgWT Vehicle (Gray bars), TgKO Vehicle (Black bars), TgWT rapamycin (Striped gray bars), and TgKO rapamycin (Striped black bars) mice at 8.5 months of age. Analysis of Saβ-gal activity (Figure C), indicated by red fluorescence (total cell number, indicated by blue fluorescence), in brain cryo-sections from WT, TgWT or TgKO mice. (Scale bar = 500 µm. *p<0.05, ∼p<0.06 for differences between genotype defined by Fishers LSD post-hoc test on a significant effect of genotype, one way ANOVA with multiple comparisons, n=8-11 per group at 8.5 months of age, data = means ± SEM)

## Discussion

The P301S mouse model overexpresses phosphorylation-prone 4R isoforms of human tau at 5-fold higher levels than endogenous murine tau in the central nervous system, leading to neuronal loss and brain atrophy that involves hippocampus and neocortex. Neurofibrillary tangle-like inclusions develop in these regions as well as in brain stem and spinal cord, leading to severe hind-limb ataxia, paralysis, an inability to feed, and death at 9 months of age due to seeding of tau in the spinal cord[44]. It was recently reported that P301S develop senescence and cognitive impairment at 6 months of age[6], followed by severe hind limb atrophy at 9 months of age[6,44]. Environmental factors can strongly modify the development of AD-like phenotypes in mouse models of AD, leading to substantial differences in the timing of emergence of AD-like deficits at different research facilities[59,60]. It is currently unknown whether this is driven by differences in diet, genetic shift, or other factors. Our previous studies where we measured the ability of various mouse models of AD to model brain cellular senescence of human disease, showed that cognition was not impaired at 6.5 months in the P301S model and that the onset of senescent cell accumulation was delayed until 10 months of age [15]. Consistent with these prior findings, the present studies did not demonstrate cognitive decline or senescence at 8.5 months of age for P301S TgWT mice. Further, we found that the onset of hind-limb paralysis was delayed to 13 months of age. It is worth considering that these delays in cognitive and motor phenotypes coincide with an absence of cellular senescence in brain, indicating a potential for senescence to be a factor in phenotypic variability of AD mouse models. However, significant progress needs to be made in our understanding of AD mouse models and senescence to establish this potential causal relationship.

In our colony, P301S mice express tau at 12-fold higher levels than endogenous murine tau, and develop hind-limb phenotype at 13 months of age. Genetic ablation of Nrf2 in P301S mice accelerated the hind-limb paralysis phenotype of this model from 13 months of age to 10 months of age, but contrary to hind-limb ataxia, loss of Nrf2 did not further increase brain levels of tau or pTau in TgKO mice compared to the TgWT group, indicating that accumulation of pTau in brain was not the trigger for hind-limb paralysis acceleration. It is possible that this discrepancy is due to the robust immunosurveillance mechanisms of the brain[61] as compared to the spinal cord. It is known that the spinal cord is highly sensitive to damage from oxidative stress compared to other tissues [62], thus depletion of Nrf2 may increase seeding of tau in the spinal cord and exacerbate the phenotype.

At 8.5 months of age, TgWT mice did not exhibit long term memory impairment or decline in cognitive flexibility in the MWM. However, in the absence of Nrf2, TgKO mice performed mildly but significantly worse than WT mice in tests of memory of platform location, indicating that loss of Nrf2 may have accelerated cognitive decline. At 8.5 months of age, TgWT and TgKO mice did not show evidence of senescent cell accumulation. While elimination of Nrf2 mildly impaired cognitive performance, it did not rely on increased senescent cell burden to do so. These data suggest that senescent cell burden in brain was not the main cause of accelerated cognitive impairment associated with Nrf2 loss. However, we did not measure senescent burden in spinal cord. It is thus possible that this region was more sensitive to loss of Nrf2, resulting in increased senescent cell burden that could be associated with hind-limb ataxia. Although cellular senescence was the focus of this study, a different group has shown that Nf2KO mice exhibit cognitive decline at 20 months of age as a result of mitochondrial dysfunction and increased synaptic dysfunction [56]. It may thus be possible that the impact of Nrf2 ablation on cognition and synaptic function relies on effects of mitochondrial dysfunction unrelated to senescent cell burden.

Lastly, a more detailed investigation on the onset of senescent cell burden in our model is needed. In our hands, P301S mice develop cellular senescence in the hippocampus at 10 months of age, before emergence of hind-limb ataxia[15]. It is possible that Nrf2 deficient mice may develop elevated levels of senescence in the brain after 8.5 months of age, but before the onset of hind-limb ataxia at 10 months of age. Since we observed a significant acceleration of hind-limb and cognitive phenotypes without an increase in senescent burden in the brain, either another tissue may play a role, or another cellular pathway may be responsible for these impairments.

DQ has been shown to eliminate senescence of aged mice in multiple tissues, including the brain, alleviating brain inflammation, bone loss, and physical dysfunction[10,63,64]. DQ treatment also alleviated NFT burden in a mouse model of tauopathy, although it was unclear whether DQ treatment had effectively cleared senescent cells in brain[65]. In the present study, long-term DQ treatment was unable to improve cognitive performance or reduce senescent burden in brains of P301S mice independent of Nrf2 status. This may be attributed to the lack of senescent cell accumulation in brains of P301S mice at this age, or alternatively DQ might be ineffective at eliminating senescent cells from the brain of this model. It has been noted that senescence is not a homogenous event, in that senescent cells from different inducers, cell types, and animal models produce unique SASP profiles as well as senescent cell anti-apoptotic pathways (SCAPs)[18]. While DQ has been shown to remove senescent cells from brains of aged mice, it has not been shown to impact senescence in brains of mouse models of AD, and had not previously been tested in the P301S model.

Rapamycin prevents senescent cell accumulation in different cell types and from multiple senescence inducers[28–30]. Numerous studies indicate that rapamycin reduces amyloid plaque burden and improves cognition in the hAPP(J20)[31–33,66], P301S(PS19)[34], 3XTg-AD[25,51,67], and viral vector-based P301L[35] mouse models of AD and tauopathy. Previous work in our lab showed that rapamycin can prevent, but not clear senescent accumulation in mouse embryonic fibroblasts as well as in the lung and adipose tissue of mice through an Nrf2 dependent manner[27]. In the present study, long term treatment with rapamycin delayed spatial learning and led to a modest decrease in spatial memory. It is important to note that the dose that was used in our studies is 4-fold higher than the dose that was used in prior studies in AD models and it is currently unknown if rapamycin administered by I.P. injection crosses the blood-brain barrier to the same extent that drug administered orally by supplementation in the chow[25,31,33,51,66,67]. If rapamycin administered via I.P. did not effectively cross the blood brain barrier, that may explain why we did not observe equal or improved cognitive performance as compared to vehicle treated mice, although spatial learning and memory in this group was indistinguishable from that of WT littermate mice. However, we did not find that rapamycin reduced senescence markers in peripheral tissue. AD mouse models also tend to be sensitive to mouse handling, so frequent I.P. injection could have possibly been a confounding factor for our results, although both rapamycin and vehicle treatments were performed by I.P. injection. Apart from being able to extend health and lifespan in various animal models, rapamycin treatment can conversely cause metabolic disorders including vascular and mitochondrial dysfunction if given at too high of a dose or for too long[68]. Because we treated our experimental groups with doses higher than those used in other prior studies in models of AD, it is possible that we have engaged some of these detrimental mechanisms although other studies using higher doses of rapamycin have shown substantial beneficial effects in a variety of models of aging and age-associated disease[49]. Taking this in consideration with the fact that rapamycin alleviates AD-like cognitive and histopathological deficits both before[32] or after disease onset[31,33] or in other models only pre-symptomatically [67], and termination of rapamycin treatment can result in the relapse of cancers[69], the use of rapamycin for AD and other dementias will likely require careful determination of adequate doses.

Ultimately, in this study we found that elimination of Nrf2 could accelerate the progression of cognitive decline in a tau-based AD model. However cellular senescence did not appear to be the mechanism through which this occurred. We found that a lack of cellular senescence coincided with a delay in phenotype development in P301S mice, indicating that cellular senescence may be a mechanism subject to phenotypic drift between mice raised at different animal facilities.

Recently, a hypothesis emerged which predicts that rather than senescence driving plaque and tangle formation in AD, senescence acts as an external mechanism for hyperphosphorylated tau and NFTs to promote neurotoxicity[65]. In postmortem human and mouse brains, it was reported that some markers of senescence are found surrounding neurons containing NFTs, but not in neurons near depositions of Aβ plaques[7,65]. Our data, which showed that a lack of senescence was associated with a delay in cognitive phenotype, and that senescence was not the mechanism through which Nrf2 loss accelerated cognitive decline, suggests that the emergence of senescence may be causally associated with onset of cognitive decline in the P301S model, and indicate that Nrf2 protects brain function through mechanisms that may include, but not require the inhibition of senescence.

## Figure Legends

**Supplementary Figure 1. No evidence of senescent burden in isolated astrocytes, PFC, or adipose tissue in P301S mice at 8.5 months of age** qPCR analysis of senescent markers from various tissue. (White bars=WT, gray bars=TgWT, black bars=TgKO, solid colors=vehicle treatment, striped colors=drug treatment) A-D) Senescent markers were not elevated in TgWT or TgKO mice in isolated whole brain astrocytes. Senotherapeutics had no effect. E-H) Senescent markers were not elevated in TgWT or TgKO mice in PFC tissue. Senotherapeutics had no effect. I-J) Senescent markers were not elevated in TgWT or TgKO mice in adipose tissue. Senotherapeutics had no effect. K-L) GFAP expression is elevated in TgWT and TgKO hippocampus compared to WT mice. (*p<0.05 for differences between genotype or treatments at 8.5 months of age, n=8-10 for astrocytes and PFC, n=4-5 for adipose tissue, one way ANOVA with multiple comparisons, data = mean ± SEM)

**Supplementary Figure 2. All genotype and treatment groups performed similarly on cued trials. Cued trials to assess swim speeds and motivation to escape maze**. A-B) No main effect of genotype for CIPL values on cued trials for Male or Female mice. C-D) No main effect of treatment for CIPL values on cued trials for rapamycin or DQ treatment

