## Supplemental Figures for "Effect of Nrf2 Loss on Senescence and Cognition of Tau-Based P301S Mice"

### Supplementary Fig. 1

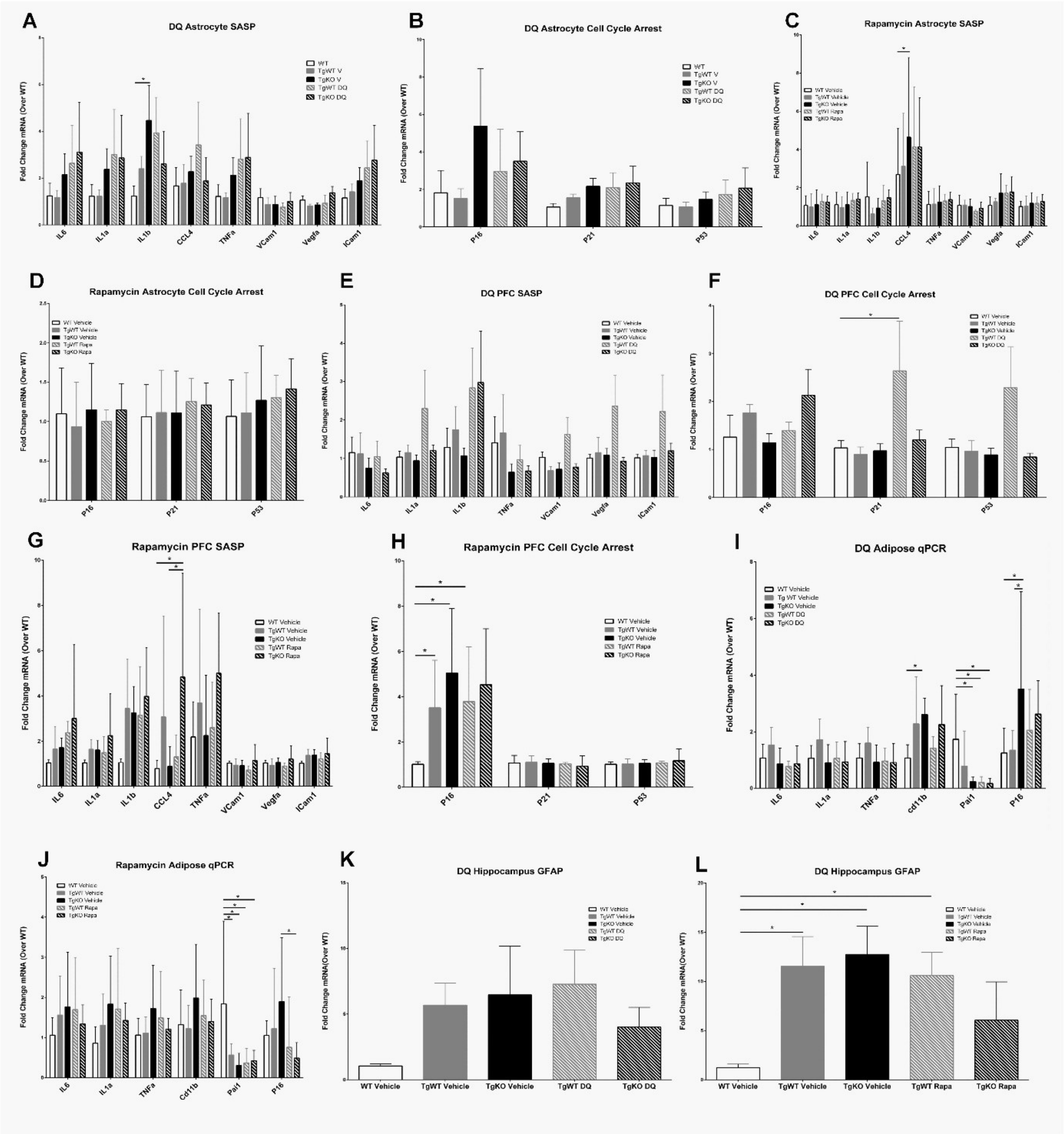

Supplementary Fig. 2

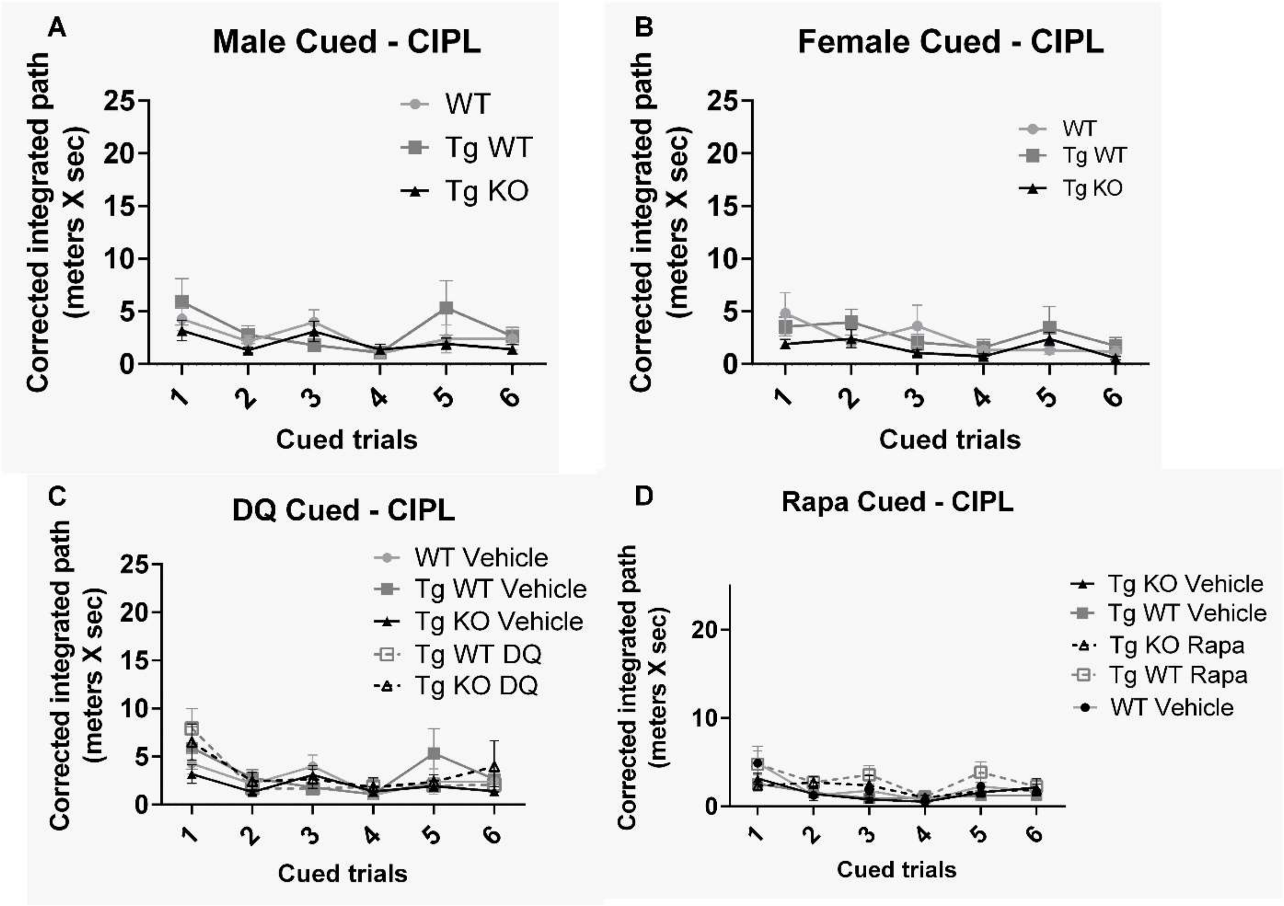

| Gene Transcript | 5' → 3' | |
|---|---|---|
| | Forward | Reverse |
| p16 | TGCAGATAGACTAGCCAGGGC | CTCGCAGTTCGAATCTGCAC |
| p21 | GTGGGTCTGACTCCAGCCC | CCTTCTCGTGAGACGCTTAC |
| p53 | CACAGCGTGGTGGTACCTTA | TCTTCTGTACGGCGGTCTCT |
| IL-6 | CCGGAGAGGAGACTTCACAG | TCCACGATTTCCCAGAGAAC |
| IL-1β | GACCTTCCAGGATGAGGACA | AGGCCACAGGTATTTTGTCG |
| TNF-α | AGCCCCCAGTCTGTATCCTT | CTCCCTTTGCAGAACTCAGG |
| PAI-1 | GACACCCTCAGCATGTTCATC | AGGGTTGCACTAAACATGTCAG |
| MCP-1 | GCTCAGCCAGATGCAGTTAA | TCTTGAGCTTGGTGACAAAAACT |
| GAPDH | ACCAACTGCTTAGCCCCCC | TGCAGGGATGATGTTCTGGG |
| β-Actin | CGCCACCAGTTCGCCATGGA | TACAGCCCGGGGAGCATCGT |
| CD11b | TCCGGTAGCATCAACAACAT | GGTGAAGTGAATCCGGAACT |
| IL1-α | CCCGTCCTTAAAGCTGTCTG | AATTGGAATCCAGGGGAAAC |
| CCL4 | CCCACTTCCTGCTGTTTCTC | GAGGAGGCCTCTCCTGAAGT |
| VCam-1 | GTGGTGCTGTGACAATGACC | ACGTCAGAACAACCGAATCC |
| ICam-1 | AGCACCTCCCCACCTACTTT | AGCTTGCACGACCCTTCTAA |
| Vegf-α | CCAGGAGGACCTTGTGTGAT | GGGAAGGGAAGATGAGGAAG |
| GFAP | AACCGCATCACCATTCCT | CGCATCTCCACAGTCTTTACC |

**Supplementary Table 1. Mouse primers** Forward and reverse primer sequences used for qPCR studies to determine senescence in mouse brain tissue
